# Quantifying data reuse in proteomics using PRIDE downloads statistics and a semi-supervised LLM-based framework

**DOI:** 10.64898/2026.04.16.718670

**Authors:** Suresh Hewapathirana, Jingwen Bai, Chakradhar Bandla, Selvakumar Kamatchinathan, Deepti J Kundu, Nithu Sara John, Boma Brown-Harry, Nandana Madhusoodanan, Joan Marc Riera Duocastella, Juan Antonio Vizcaíno, Yasset Perez-Riverol

**Affiliations:** European Molecular Biology Laboratory, European Bioinformatics Institute (EMBL-EBI), Wellcome Trust Genome Campus, Hinxton, Cambridge CB10 1SD, UK

**Keywords:** download hubs, data reuse, download analytics, LLM-refined classification, PRIDE, proteomics

## Abstract

Understanding how scientific datasets are accessed and reused is essential for resource planning and impact assessment. Here we present the PRIDE Archive download tracking infrastructure and a comprehensive analysis of 159.3 million download records from the PRIDE proteomics database (2021-2025), spanning 35,528 datasets accessed from 235 locations. The infrastructure includes nf-downloadstats, a scalable Nextflow pipeline for processing download logs, and DeepLogBot, a machine-learning framework that classifies traffic into bots, institutional download hubs, and independent user downloads. DeepLogBot combines heuristic seed selection with multi-LLM annotation (Claude and Qwen3) to produce gold-standard training labels, achieving 92.2% bot classification accuracy on a held-out test set. After separating bot traffic, analysis reveals downloads from 214 countries/regions, 249 institutional download hubs, and a concentrated reuse distribution, with the top five countries (United States, United Kingdom, Germany, China, and Canada) accounting for over 54% of independent user downloads. These findings provide actionable insights for repository infrastructure planning and highlight the importance of distinguishing automated from individual access in scientific data resources.

## Introduction

The PRIDE database is the world-leading data repository for mass spectrometry-based proteomics [1]. As a founding member of the ProteomeXchange consortium [2], PRIDE enables researchers to share and access proteomics datasets globally, promoting transparency, reproducibility, and data reuse. Aligned with the FAIR principles - Findable, Accessible, Interoperable, and Reusable [3] - PRIDE supports open science by ensuring that public datasets are well-annotated and machine-readable. These principles are essential for maximizing the value of shared scientific data.

Understanding how much public datasets are reused is essential for assessing the scientific impact of a data resource such as PRIDE and informing data-driven policies. While citations in scholarly publications offer one indicator of reuse, data download statistics can provide a more granular view of data demand from any data resource. In our previous work [4], we demonstrated that usage/downloads metrics can serve as complementary indicators of scientific impact for a given publication, supporting improved data stewardship, resource allocation, and funding decisions. Beyond measuring the global impact of a data resource, download statistics are critical for designing more effective data infrastructures. Download patterns can inform the optimization of data access protocols, guide the prioritization of metadata curation and visualization features, and identify high-value datasets for targeted annotation or integration efforts. As public data volumes continue to grow, usage-driven strategies become increasingly important for improving dataset discoverability and reuse. For example, in the concrete case of PRIDE, frequently downloaded datasets - particularly those used as community benchmarks - could be prioritized for enhanced manual metadata curation/annotation (e.g., SDRF-based sample metadata [5]) and enriched with curated tags and keywords, making them easier to find through search interfaces. Users could then combine these annotations with download counts as a proxy to identify the most relevant and community-validated datasets for reuse. Similarly, repositories can leverage download patterns to allocate faster transfer services and optimized storage for high-demand datasets, ensuring that the most reused data remains readily accessible. In contrast, knowing which datasets are less frequently downloaded can help repositories to identify underused datasets that may benefit from improved metadata and file organization.

Despite their importance, systematic tracking of dataset downloads remains a major challenge across bioinformatics resources in general, including PRIDE and the other ProteomeXchange partners. Despite some common access statistics having been widely adopted across data resources [6], barriers include the absence of a standardized infrastructure for logging access events, technical complexities in aggregating usage data across distributed and heterogeneous transfer systems (e.g., FTP, HTTP), and ongoing concerns related to user privacy and data protection. Compounding these challenges, automated bot traffic contaminates download statistics - studies estimate that bots account for 30-70% of all internet traffic (https://cpl.thalesgroup.com/ppc/application-security/bad-bot-report), and scientific repositories are particularly attractive targets due to their open-access policies and valuable content [7]. Without accounting for this contamination, any analysis of repository usage risks will conclude inflated and distorted metrics. As bioinformatics resources continue to scale in both size and complexity, robust download analytics will become increasingly vital - not only for measuring impact - but also for enabling smarter, user-informed development of open data platforms.

Here, we present the PRIDE Archive download tracking open infrastructure, which includes nf-downloadstats, a large-scale Nextflow [8] workflow for processing extensive traffic logs, and DeepLogBot, a machine-learning framework that classifies download traffic into bots, institutional download hubs, and independent user access. Additionally, we have developed an infrastructure that integrates download statistics directly into the PRIDE Archive web interface, allowing users to sort datasets (e.g., as the result of searches) by total downloads - both raw and normalized by number of files in the dataset - and perform percentile-based analyses. Using these tools on 159.3 million download records spanning 2021 to 2025, we characterize global proteomics data download patterns as a proxy for data reuse, examining geographic distribution across countries and regions, temporal evolution of download activity, shifts in download protocol preferences, and the concentration of dataset downloads across the PRIDE collection.

The workflow is open-source, and we make available the resulting data, which can be used by PRIDE data submitters, e.g., in grant reports and publications.

## Materials and Methods

### nf-downloadstats: Log Processing Pipeline

Download logs are stored in the EBI (European Bioinformatics Institute) file system as compressed, comma-delimited text files. Each log entry includes a timestamp, dataset accession, filename, anonymized and non-reversible IP hash, download status, geographic location (geo), used download protocol (Globus, HTTP, Aspera, FTP), and dataset type. No personal or directly identifiable user data is stored. Log files are organized hierarchically by Protocol, Public/Private access, Year, Month, and Day. Individual log files can be large, ranging from 1 GB to 237 GB (**Supplementary Note 1**).

Due to the large volume and size of download log files, we developed nf-downloadstats (https://github.com/PRIDE-Archive/nf-downloadstats), an open-source Nextflow workflow for large-scale processing of the original anonymized log files (**Figure 1**). Each processing step is implemented as an independent Python module, performing tasks such as removing incomplete transfers, removing log inconsistencies, and selecting PRIDE-specific records. To efficiently process the large volume of log files, parallelization is employed using a high-performance computing (HPC) environment managed via Slurm. Log files are processed in batches, with batch sizes and filtering criteria defined in a user-friendly YAML configuration file. The output is a consolidated 4.7 GB Parquet file containing 159,327,635 individual download records spanning five years: from January 2021 through December 2025 (2025 data extends to December 10). Each record includes the download date, geolocation, dataset accession, filename, and download method (protocol). We aggregate download events at the geolocation level (*geo_location*), carrying forward the associated *country* and *geoip_city_name* labels from the traffic logs; because some records originate from proxies, cloud endpoints, disputed or overseas territories, or non-standard geocoding labels, not all locations can be unambiguously mapped to a single sovereign country; we will continue mentioning within the manuscript country/regions. For analysis, the full 159.3 million download events are aggregated at the *location* level, where each location represents a unique geographic coordinate. Throughout this work, a “unique session” is defined as a distinct anonymized IP hash; because IP addresses are hashed irreversibly at ingestion, we cannot link users across locations or identify individuals but can count distinct downloaders per location. Each location profile is characterized by behavioral features including download volumes, user counts, temporal patterns (*working hours ratio, night activity, hourly entropy*), protocol usage (HTTP, FTP, Aspera, Globus shares), burst patterns, user coordination scores, etc. Full feature descriptions are provided in **Supplementary Note 2**.

**Figure 1:**
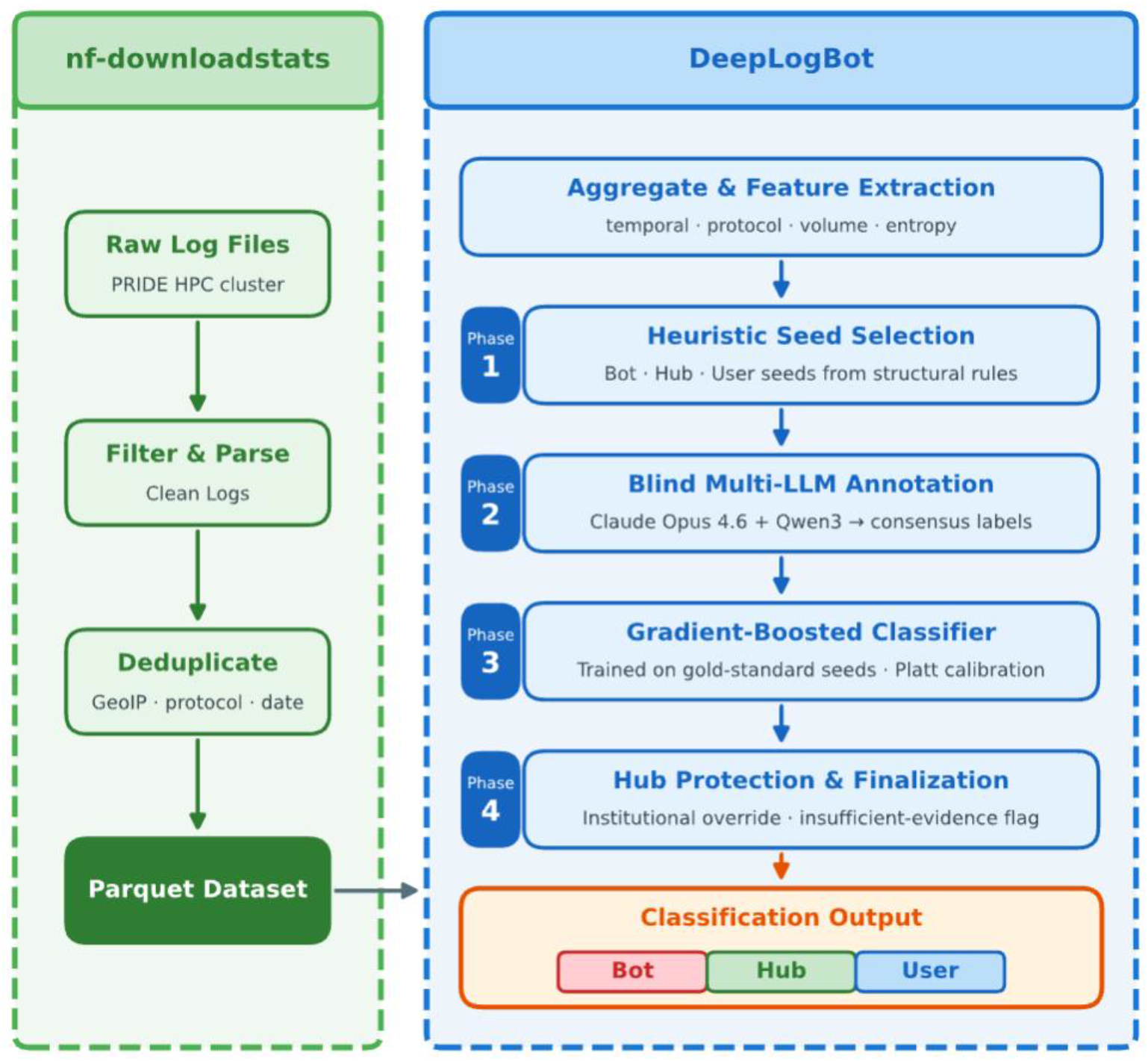
PRIDE Download Traffic Classification Pipeline. End-to-end pipeline comprising two components: nf-downloadstats (green dashed box) processes raw PRIDE download logs into a consolidated Parquet file (159M records, 4.7 GB), and DeepLogBot (blue dashed box) aggregates records to geographic locations, extracts behavioral features per location, and applies a four-phase classification pipeline (heuristic seed selection, blind multi-LLM seed refinement, gradient-boosted fusion meta-learner with Platt calibration, and hub protection with finalization) to classify locations as bot, hub, or user.

### Traffic Classification Framework

To distinguish automated traffic from individual user access, we developed DeepLogBot, a semi-supervised four-phase classification pipeline (**Figure 1**). Each geographic location is classified as a bot, hub (legitimate automation), or user. In Phase 1, heuristic rules derive training seeds from structural signals: user seeds via a three-tier system (individual researchers, active researchers, research groups), bot seeds via six complementary signals (bot-farm, distributed, nocturnal, coordinated, scraper, and explosive-growth patterns), and hub seeds from institutional mirrors and sustained high-volume sites. In Phase 2, to break the circularity of heuristic-only training, 1,153 locations sampled across 20 stratified feature-space zones were annotated blindly by two LLMs, Claude Opus 4.6 and Qwen3-30B-A3B, using only behavioral features and geographic context, yielding 934 consensus labels. These were split into training (67%) and held-out test (33%) sets and injected as high-confidence seeds that override heuristic labels. In Phase 3, a gradient-boosted classifier (200 estimators, max depth 5) is trained on the gold-standard labels using 36 behavioral features. Phase 4 applies post-classification hub protection: locations with institutional download patterns (specialized protocols, multi-year activity) are never classified as bots; flags locations with fewer than 3 downloads as insufficient evidence and derives final Boolean labels. Full pipeline details are in **Supplementary Notes 3-6**.

Training on gold-standard consensus labels improved classification accuracy from 62.1% (heuristic seeds only) to 92.2% on the held-out test set of 309 locations. The largest gains came from correctly reclassifying research-city locations previously mislabeled as bots and residential-area bots previously mislabeled as independent users. Bot locations are typically characterized by large numbers of anonymized users each making very few downloads (3-15 downloads per user), uniform temporal patterns lacking circadian rhythm, and activity concentrated in a single year, consistent with distributed web crawlers, scrapers, or automated scanning tools.

## Results

### Global PRIDE Usage Patterns

Over the 2021–2025 study period, 159.3 million individual file downloads were recorded across 35,528 datasets, accessed from 235 countries. After applying our classification pipeline (**Figure 1**), 48.2% of traffic was identified as originating from 27,063 automated bot locations and separated from hub traffic and independent user access. The remaining 249 institutional download hubs (50.0% of total traffic) and independent user downloads reveal the true scale of researcher engagement with PRIDE data (**Figure 2A**).

**Figure 2:**
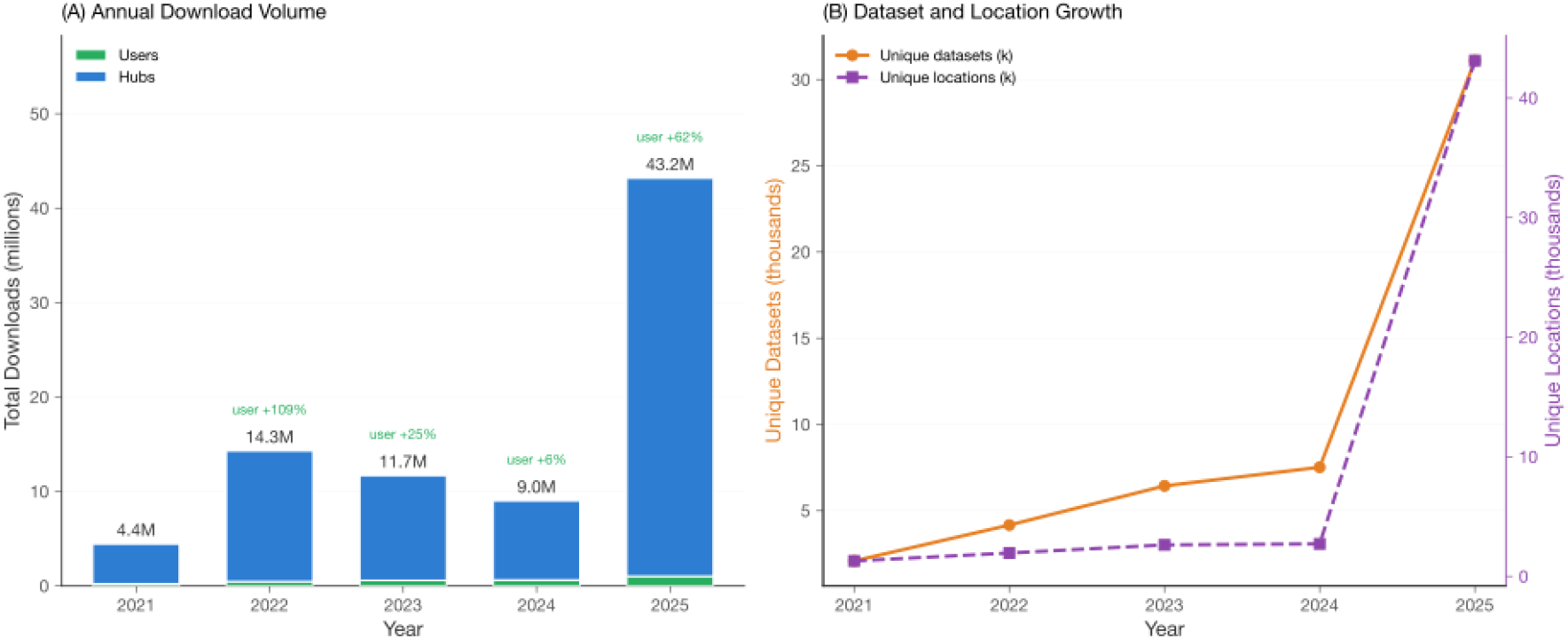
Temporal trends in PRIDE usage. (**A**) Annual download volumes for users and hubs, with year-over-year user growth percentages annotated. (**B**) Growth in unique datasets accessed and unique geographic locations over time.

The geographic reach of PRIDE data reuse is truly global. After separating bot and hub traffic, user downloads span 214 countries/regions with broad geographic diversity. To characterize the relationship between user base size and download intensity, we plotted user-only downloads against unique users for the top 50 countries (**Figure 3A**). Download patterns vary considerably: some countries show broad user bases with moderate per-user activity, suggesting predominantly individual researchers, while others exhibit high per-user averages reflecting concentrated institutional access.

**Figure 3:**
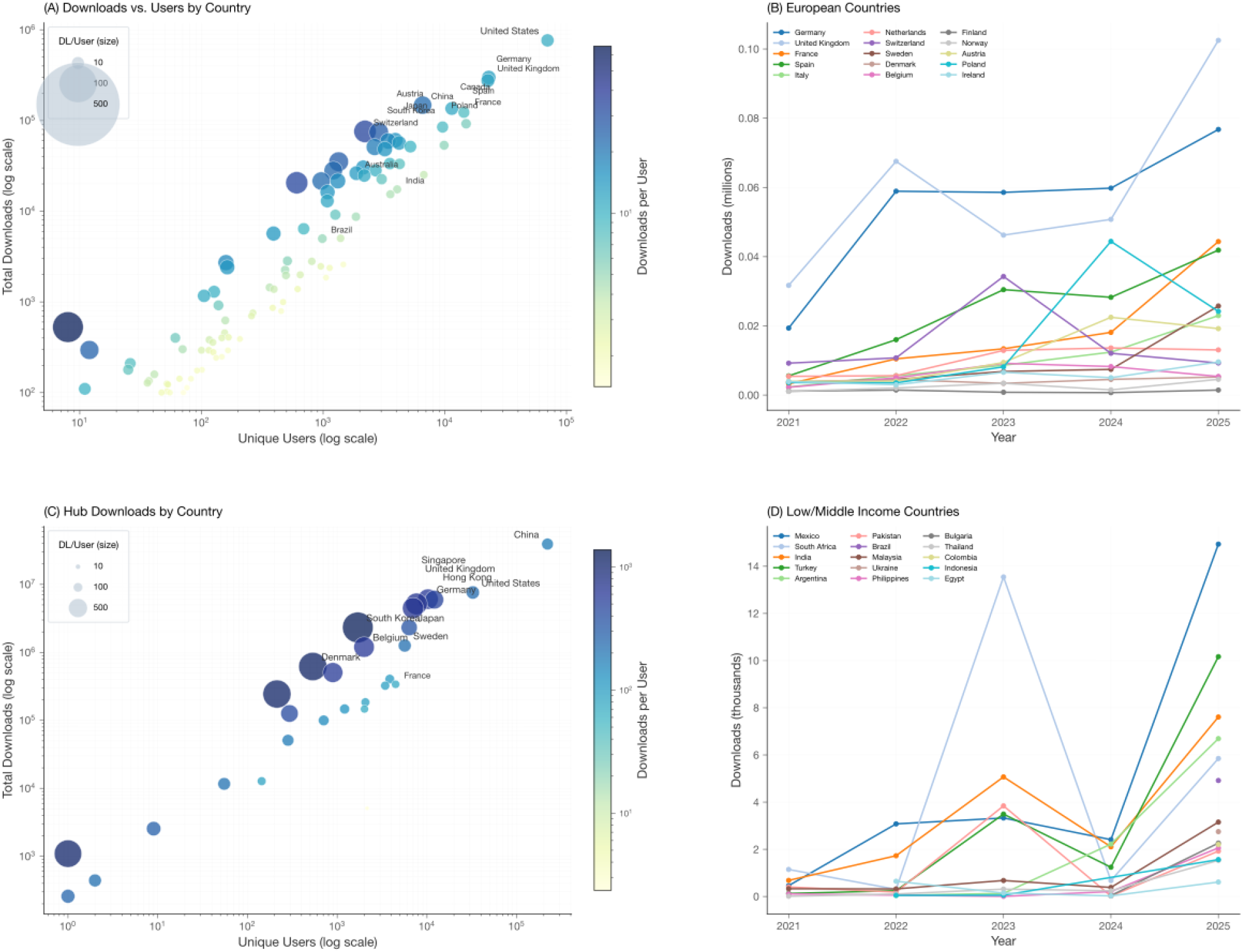
Global country-level download patterns. (**A**) User downloads versus unique users for the top 50 countries (bot and hub-filtered); bubble size and color represent downloads per user on a log scale. (**B**) Yearly trends for the top 15 European countries. (**C**) Hub downloads by country. (**D**) Yearly trends for low- and middle-income countries.

Although European countries account for a major share of user downloads, yearly trends reveal shifting dynamics (**Figure 3B**). Notably, PRIDE usage is growing in some low- and middle-income countries (LMIC, as defined by the Wellcome Trust based on the OECD DAC list (Wellcome Trust 2025); **Figure 3D**), suggesting that PRIDE is increasingly serving as a resource for researchers in developing nations, supporting broader global participation in proteomics data reuse.

### Download Concentration

Dataset user downloads (after bot and hub removal) follow a highly skewed distribution characteristic of heavy-tailed systems (**Figure 4A**). The rank-frequency distribution reveals a characteristic long tail, with download counts dropping steeply beyond the top datasets. A ranking of the top 20 most downloaded datasets (**Figure 4B**) shows that community benchmark resources and tissue atlas datasets dominate. The most downloaded dataset is **PXD010154** (31.4K user downloads from 43 countries), while the ProteomeTools synthetic peptide libraries (**PXD004732**, ranked 8th) reflect widespread use as training data for machine learning models and spectral library search engines. Importantly, the most downloaded datasets are not one-time events but show sustained behavior over multiple years (**Figure 4C**). Of the top 25 datasets, most have been actively downloaded in at least 4 of the 5 years covered (2021–2025). Extended analyses are provided in **Supplementary Note 7**.

**Figure 4:**
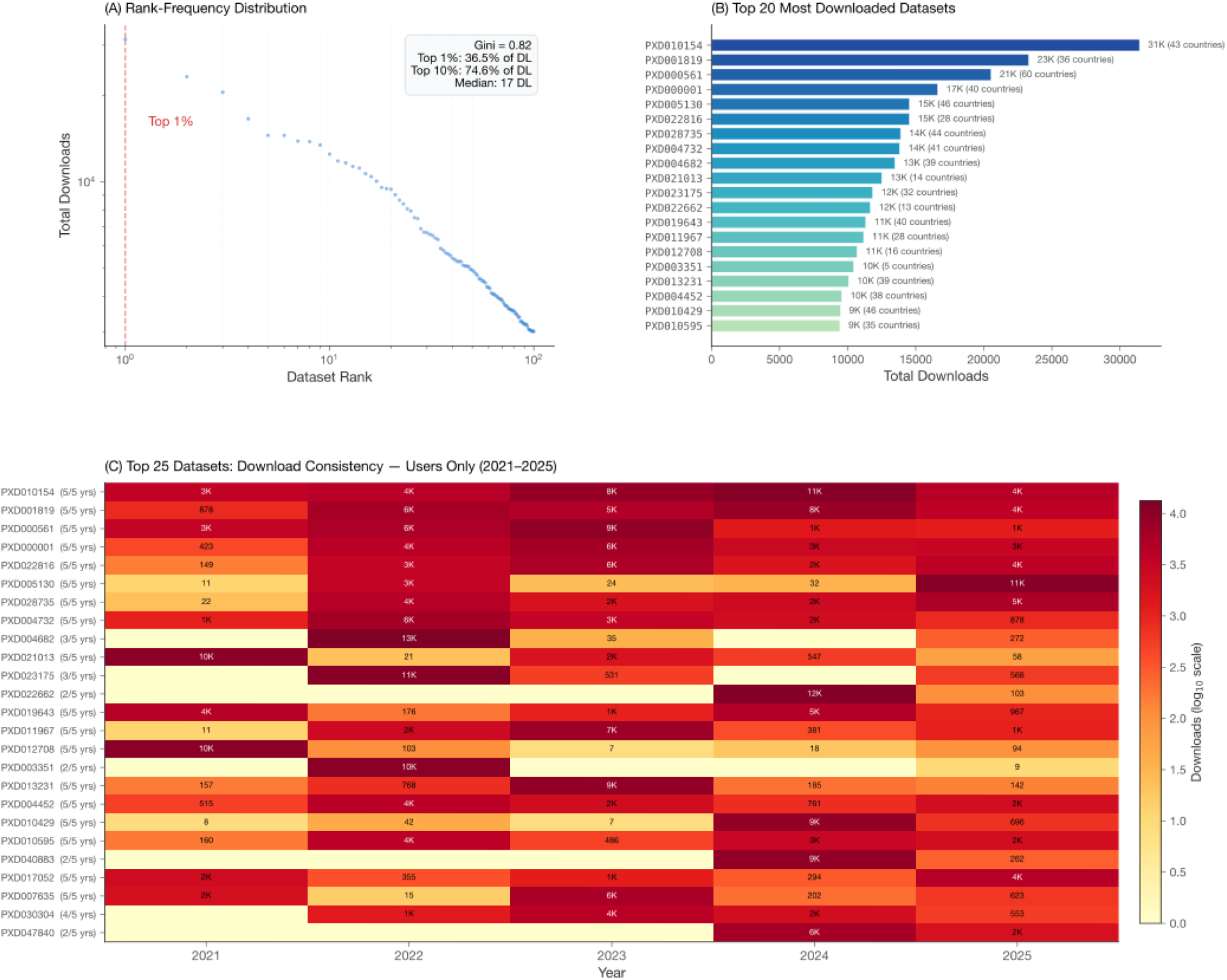
Dataset download concentration and consistency (user downloads only, after bot and hub removal). (**A**) Rank-frequency distribution on a log scale; the dashed red line marks the top 1% boundary. (**B**) Top 20 most downloaded datasets with the number of accessing countries. (**C**) Download consistency heatmap for the top 25 datasets (2021–2025); color intensity represents download count on a log_10_ scale; most top datasets show sustained volumes across 4-5 years, indicating their role as community reference and benchmark datasets.

Download activity, particularly sustained, multi-year download patterns, provides a complementary measure of genuine community adoption that is not fully captured by publication records alone. By separating bot and hub traffic from individual user downloads, the download statistics presented here more accurately reflect individual researcher engagement with specific datasets. We want to emphasize that some of these values; particularly small, granular differences between datasets, such as shifts in ranking positions; may be influenced by limitations in the bot-detection algorithm (**Supplementary Note 8**). While we are confident that the algorithms capture the overall download patterns robustly, finer differences across datasets should be interpreted with caution, as they may vary substantially.

### File Transfer Protocol Usage

Before 2025, PRIDE downloads relied almost exclusively on HTTP and FTP, with FTP dominating in 2021 (66.7%) and 2023 (61.2%), and HTTP leading in 2022 (73.4%) and 2024 (60.2%) (**Figure 5A**). This shifted in 2025 with the emergence of Aspera (FASP), which accounted for 10.5% of all non-bot downloads (4.5M downloads) despite appearing only from July onward (**Figure 5B**). Aspera usage peaked in September 2025 with 3.2M downloads, driven primarily by institutional hubs in China, particularly a hub in Chongqing (5.1M total downloads, 70% via Aspera), Hefei (182K downloads, 93% via Aspera), and a hub in Kensington, MD near NIH/NCBI (36K downloads, 11% via Aspera). The timing of Aspera adoption coincides with the release of pridepy [9] in March 2025, a Python-based command-line tool that abstracts protocol complexity and enables seamless switching between FTP, Aspera, and Globus transfers with a single command, substantially lowering adoption barriers for high-performance protocols (**Supplementary Note 7**).

**Figure 5:**
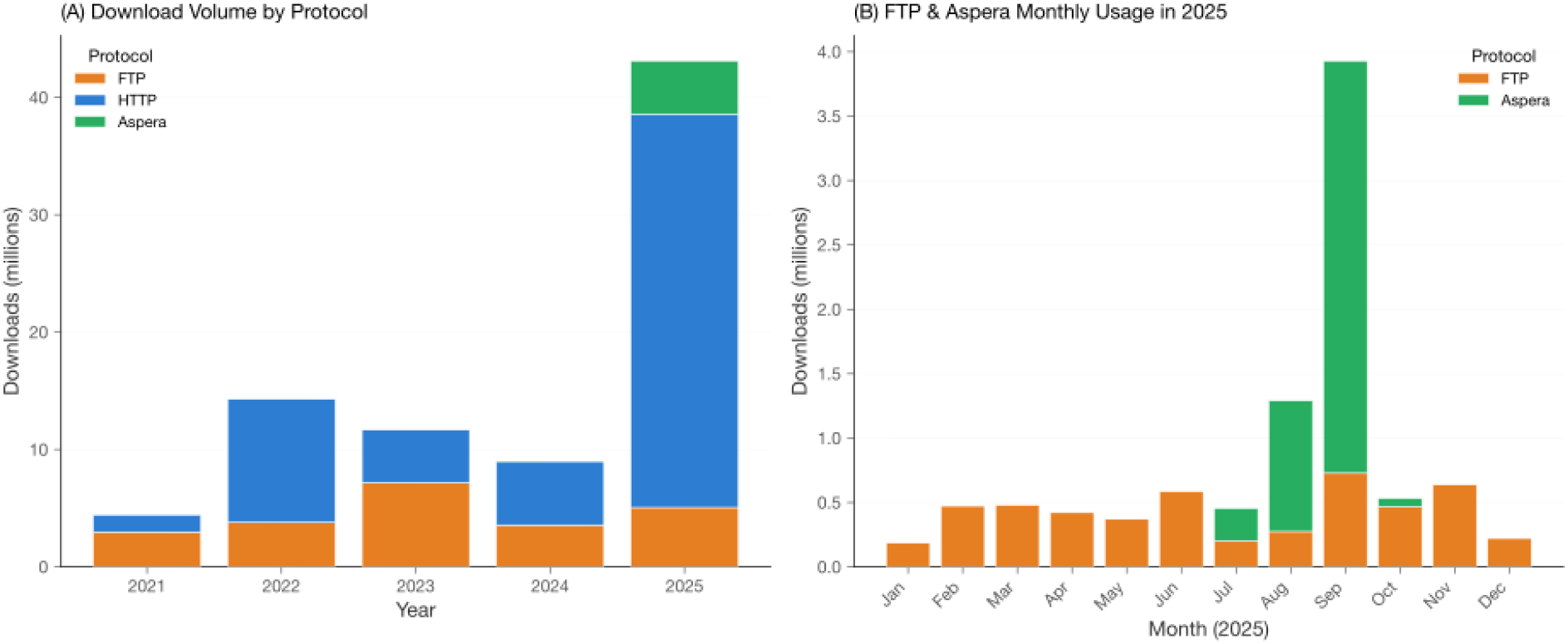
Protocol usage for independent user and hub downloads (after bot removal). (A) Annual download volume by protocol. (B) Monthly FTP and Aspera usage in 2025, showing the emergence of high-performance transfer protocols.

### Download Hubs

Institutional download hubs are characterized by few users with very high per-user download rates, usage of specialized bulk-transfer protocols (Aspera, Globus, FTP), and sustained multi-year activity (**Figure 6**). These hubs represent institutions that systematically and continuously download public proteomics data for reanalysis, mirroring, or aggregation purposes. The geographic spread of hubs, spanning all six inhabited continents, demonstrates that institutional data reuse is a global phenomenon. Hub characteristics vary widely (**Figure 3C**): some operate with very few users but extremely high per-user download rates, consistent with mirrors or automated reanalysis pipelines, while others involve many users accessing data at moderate intensity, suggesting shared institutional infrastructure.

**Figure 6:**
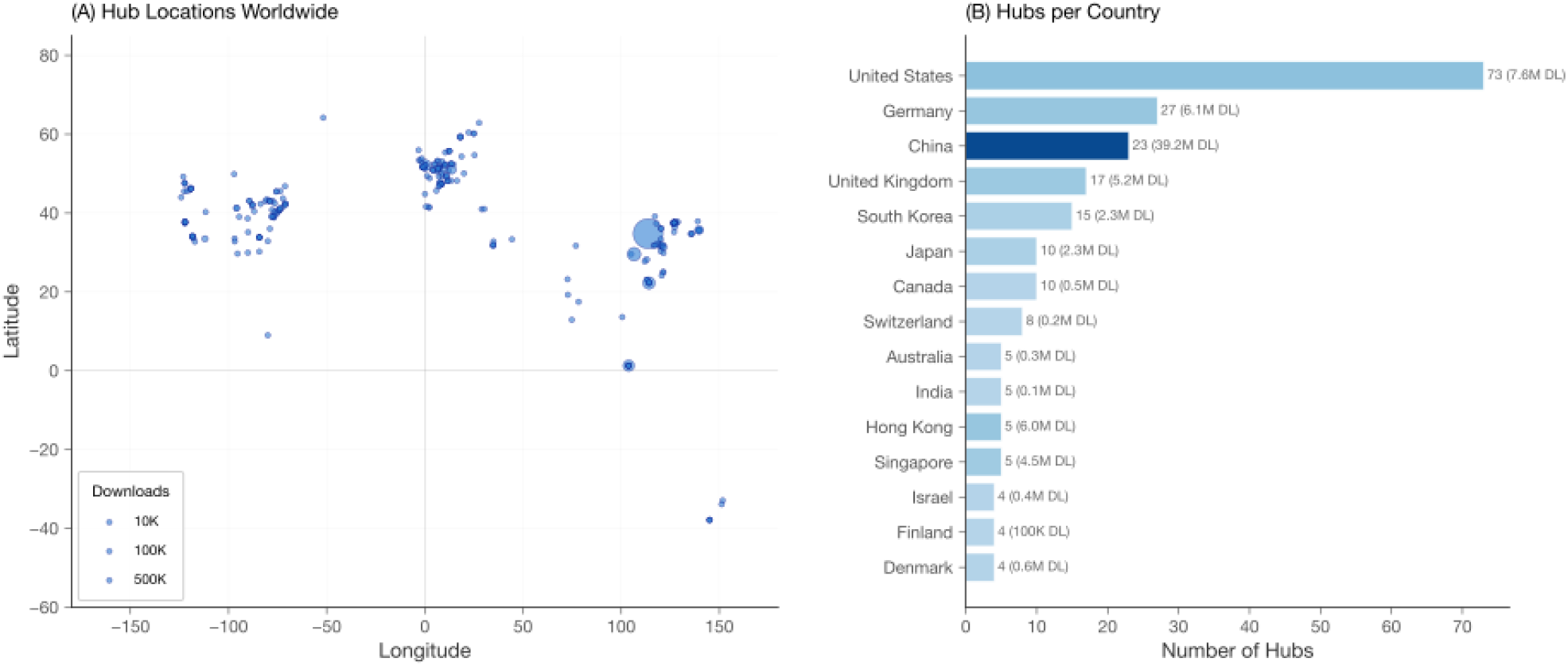
Distribution of 249 identified download hubs. (A) Geographic locations worldwide, with bubble size proportional to download volume. (B) Top 15 countries by hub count, annotated with total hub download volume.

### File Type Download Patterns

Analysis of file type download patterns across regions reveals distinct usage profiles (**Figure 7**): raw instrument files dominate downloads in all regions, accounting for 72–73% of traffic in East Asia and North America. LMIC countries show a lower raw file proportion (54%) with a corresponding increase in result files and processed spectra (peak list files). This imbalance highlights that most users currently need to download and reprocess raw data from scratch, even when search engine results already exist within the submission - underscoring the need for better infrastructure to make analysis results more discoverable and independently downloadable.

**Figure 7:**
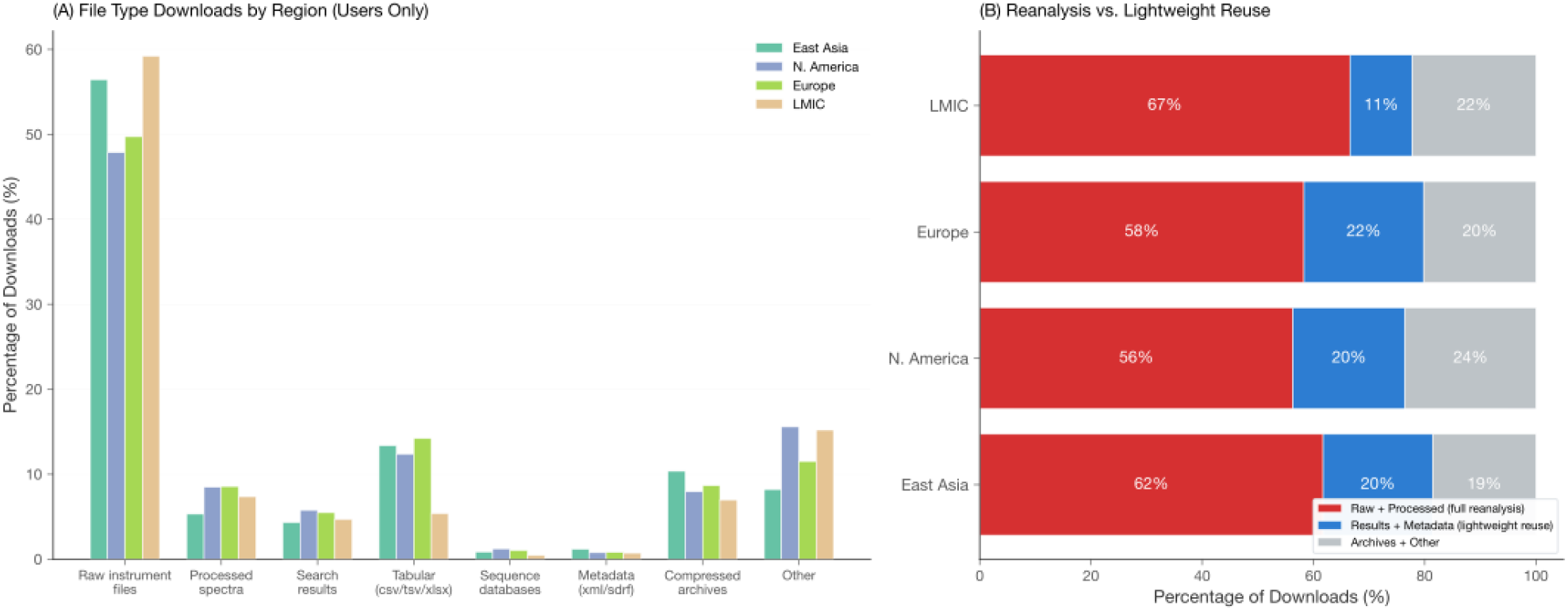
File type download patterns by region (independent user downloads only, after bot and hub removal). (A) Percentage of downloads by file type category across four regions. LMIC countries show lower raw file usage and higher result/processed spectra downloads compared to East Asia and North America. (B) Aggregated view: raw + processed spectra (full reanalysis capability) versus results + metadata (lightweight reuse). LMIC countries have the lowest reanalysis-oriented profile (55%) and the highest lightweight reuse proportion (15%).

### PRIDE Download Statistics Visualization

The aggregated download statistics produced by nf-downloadstats are stored in MongoDB and Elasticsearch to enable fast searching and visualization within the PRIDE Archive web interface. For each dataset, the total number of downloads is displayed alongside a gradient bar indicating its download percentile, with higher intensity representing datasets in the top 1% of downloads (**Figure 8**). Additionally, a yearly trend chart is provided for each dataset, illustrating download activity over time. Datasets can be sorted by both total downloads per project and a normalized metric that accounts for the number of files within each project, enabling users to identify highly reused datasets for benchmarking or reanalysis. These features allow PRIDE data submitters to use download statistics in grant reports and publications and provide the community with a tool to discover highly reused datasets.

**Figure 8:**
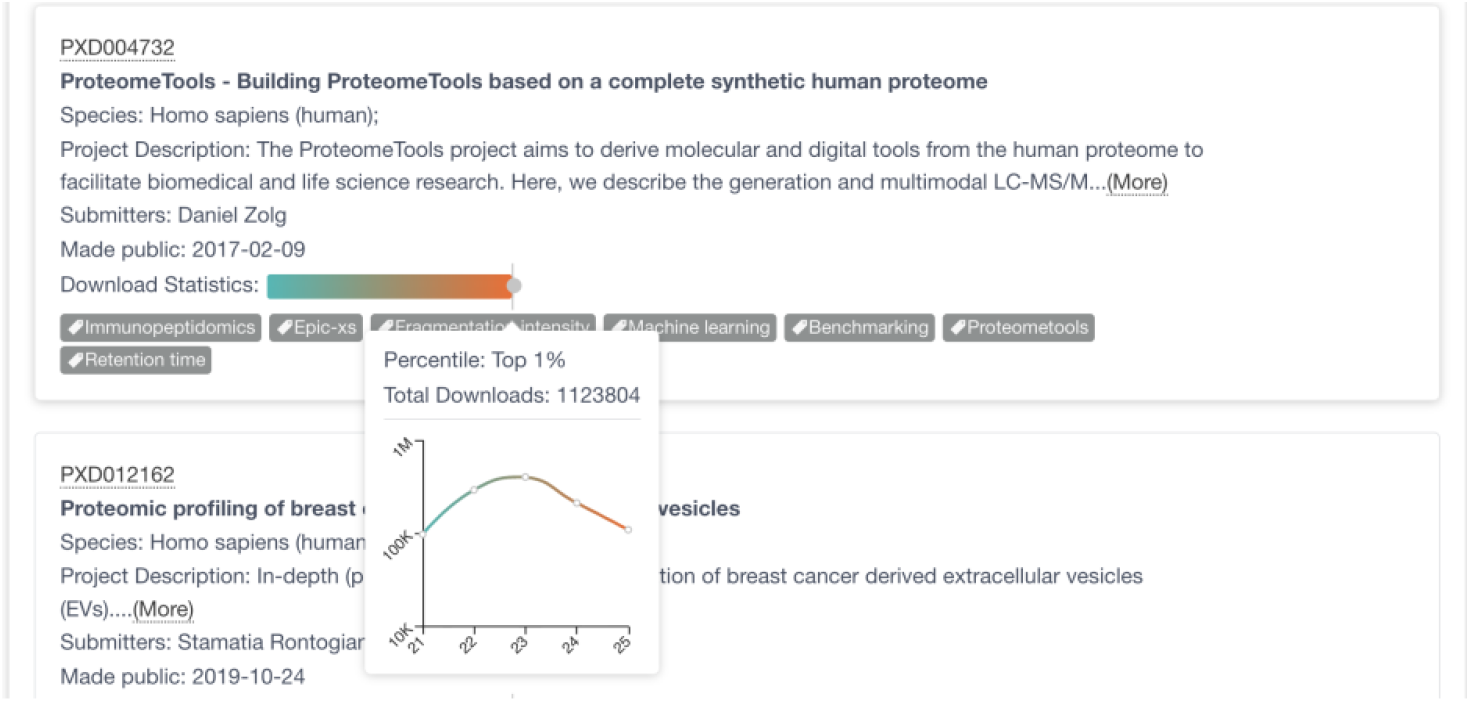
PRIDE Archive download statistics integration. Each dataset displays a gradient percentile bar and total download count. A pop-up trend chart shows yearly download activity, enabling users to assess dataset popularity and reuse trends directly within the PRIDE interface.

## Discussion

Here we have performed a detailed study of the PRIDE data download statistics for the last 5 years (2021–2025). A central finding is that 48.2% of PRIDE download traffic originates from 27,063 automated bot locations. After removing bot traffic, the remaining downloads comprise 249 institutional download hubs (50.0% of total traffic) and independent user downloads, together representing the valuable, legitimate use of PRIDE data. Without separating bot traffic, raw download volumes are unreliable as scientific impact indicators [4]. As AI-driven platforms that perform large-scale automated reanalysis become more prevalent, repositories will need adaptive classification schemes that evolve alongside legitimate automation patterns.

PRIDE download volumes have grown substantially over 5 years, confirming accelerating data access across a geographically broad user base (214 countries/regions). Download intensity varies around the world: some countries exhibit broad individual user bases, while others show concentrated institutional access, suggesting that the nature of data downloads, individual exploration versus systematic reanalysis, differs between research communities. The 249 download hubs we identified reveal a global infrastructure of institutional data consumers, from single-user mirrors performing full-repository synchronization to multi-user reanalysis centers processing hundreds of datasets. This hub distribution provides an empirical map of where proteomics bioinformatics infrastructure exists and can inform ProteomeXchange decisions about potential PRIDE mirrors placement, edge caching, and regional resource allocation; for instance, countries with growing user bases but no local hubs may benefit from targeted infrastructure support. Protocol analysis reveals that hub traffic is strongly characterized by Aspera, FTP, and Globus usage, while individual users predominantly rely on HTTP. Tools such as pridepy [9] should lower adoption barriers for high-performance protocols as datasets continue to grow in size.

User download activity grew steadily over the study period, with year-over-year increases of 109% (2022), 25% (2023), 6% (2024), and 62% (2025), representing a genuine expansion of the PRIDE user base. The 2025 surge in total download volume is largely driven by a rapid expansion of institutional download hubs, particularly in China: new hubs in Chongqing, Changsha, Wuxi, Nantong, and Lu’an, each contributing 99-100% of their traffic in 2025 alone, account for much of the hub growth. Chinese hubs collectively contributed 22.4 million downloads in 2025, representing 53% of all hub traffic that year. This expansion reflects growing investment in proteomics data infrastructure [10, 11] and the broader trend of regional data mirroring to serve local research communities, consistent with the growth of platforms such as iProX [12] and the National Genomics Data Center. The absence of a centralized PRIDE mirror in the region likely contributes to this pattern: without a local mirror, individual institutions in major proteomics cities independently download and cache large portions of PRIDE data, as the slow cross-continental transfer speeds make it more practical to pre-download datasets before they are needed for analysis. The emergence of these hubs highlights the value of establishing formal mirror agreements to coordinate replication, reduce redundant transfers, and improve data access latency for the growing Chinese proteomics community.

Dataset user downloads (after bot and hub removal) show a concentrated distribution. While community reference datasets such as ProteomeTools (**PXD004732**) show sustained multi-year downloads, likely because its comprehensive synthetic peptide spectral libraries serve as training data for machine learning models, retention time predictors, and spectral library search engines across the field, the “long tail” of rarely downloaded datasets should not be disregarded: these datasets may gain future value through meta-analyses, machine learning applications, or integration into multi-omics studies. Repositories can better serve both ends of this distribution by investing in improved discoverability (richer metadata, curated tags, and recommendation systems) alongside prioritized access for high-demand datasets.

Regional differences in file type usage, with LMIC countries showing higher reliance on processed results rather than raw files, suggest that computational capacity and bandwidth constraints shape data download patterns. The dominance of raw file downloads across all regions (**Figure 7A**) indicates that researchers currently lack easy access to analysis results within submissions, forcing them to re-download and reprocess raw data even when search engine outputs already exist. To address this, the PRIDE team is developing dedicated infrastructure for discovering, browsing, and downloading result and analysis files independently of the full raw dataset, enabling researchers with limited computational resources to directly access quantification tables, identification lists, and processed spectra (peak list files) without the overhead of re-running search engines. In parallel, the PRIDE team will prioritize SDRF sample metadata annotation [5] for the most downloaded and community-relevant datasets identified in this study, making these high-impact submissions immediately reusable through standardized experimental design descriptions. Furthermore, the PRIDE team is actively pursuing collaborations and training initiatives with research groups in low- and middle-income countries to promote the benefits of proteomics data reuse. Combining locally generated proteomics data with publicly available datasets from PRIDE can accelerate research in these regions without requiring data generation for every study. Such integrative approaches could also catalyze the establishment of local download hubs in LMIC countries, strengthening regional bioinformatics infrastructure and reducing dependence on cross-continental data transfers. Several complementary efforts support this vision: quantms (https://quantms.org) [13] generates standardized reanalysis outputs from public datasets, the PTMExchange initiative (https://www.proteomexchange.org/ptmexchange) provides harmonised results coming from the reanalysis of PTM-enriched datasets, and the PRIDE team is collaborating with developers of widely used search engines, including DIA-NN, MaxQuant, and MSFragger, to define standardized submission guidelines that ensure result files, quantification tables, and metadata are structured for immediate reuse.

In summary, we present the PRIDE database download tracking infrastructure, comprising nf-downloadstats and DeepLogBot, and the first comprehensive analysis of PRIDE data download statistics, processing 159.3 million records spanning 2021-2025. Our classification pipeline separates bot traffic, institutional download hubs, and independent user access with 92.2% accuracy on a held-out test set. After removing automated traffic, the remaining legitimate downloads across 35,528 datasets and 214 countries reveal a globally distributed user base, shifting protocol preferences, a concentrated download distribution, and diverse patterns of data reuse. The PRIDE team has integrated download statistics into the PRIDE web interface, enabling data submitters to use these metrics in grant reports and publications. Through pridepy [9] and dedicated infrastructure for result-level data access, we aim to lower barriers for researchers, particularly in LMIC, to discover and download analysis outputs without reprocessing full raw datasets.

## Data and Code Availability

The nf-downloadstats pipeline is available at https://github.com/PRIDE-Archive/nf-downloadstats and the DeepLogBot software at https://github.com/pride-archive/deeplogbot, both under the Apache 2.0 license. Download log data is available upon request from the PRIDE team.

## Author contributions

S.H. implemented the Nextflow workflow and collected the data; J.B. implemented the web interface for the download statistics components; C.B. and S.K. implemented the integration of the statistics components in the backend of PRIDE and databases; D.J.K., N.S.J., B.B-H., and N.M. contributed to reviewing the manuscript, the data generated, and curated some of the datasets; M.R.D. generated the infrastructure for log anonymization and provided the log files to the PRIDE team; J.A.V. reviewed the manuscript; Y.P-R. designed the study, developed the bot detection framework, performed the analysis, and wrote the manuscript.

## Acknowledgements

This work was supported by EMBL core funding; Wellcome [223745/Z/21/Z; 301300/Z/24/Z]; Biotechnology and Biological Sciences Research Council [BB/Y513829/1, BB/S01781X/1, BB/V018779/1, BB/X001911/1]; Engineering and Physical Sciences Research Council [EP/Y035984/1]; UK Research and Innovation [UKRI701]; US National Science Foundation [NSF/2324278]. We thank the PRIDE team for their support and feedback on the development of the download tracking infrastructure and analysis. We also thank Professor Bernhard Kuster for the original discussion about this topic in 2024 during the HUPO conference in Dresden.

## Conflict of Interest

The authors declare no conflict of interest.

## Data Availability Statement

All the post-processed data and statistics are present in the following repository in GitHub https://github.com/pride-archive/deeplogbot, both under the Apache 2.0 license. Download log data big data from PRIDE is available upon request from the PRIDE team.

## Notes

### Competing Interest Statement

The authors have declared no competing interest.

