## Supplementary Notes for "Quantifying data reuse in proteomics using PRIDE downloads statistics and a semi-supervised LLM-based framework"

### **Supplementary Note 1: Data Processing Pipeline**

**Log Processing Workflow:** PRIDE download logs are processed through the nf-downloadstats Nextflow pipeline (**Figure** **1**), which retrieves raw TSV log files, parses and filters download events, merges records into a consolidated Parquet file, and generates statistics reports. The pipeline produces a 4.7 GB Parquet file containing 159.3 million records optimized for columnar analytics.

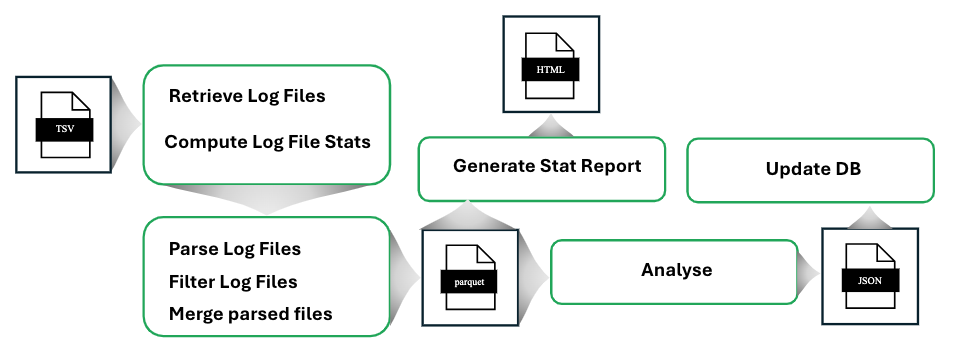

**Figure 1**: The nf-downloadstats workflow for processing PRIDE download logs. Raw TSV logs are parsed, filtered, and merged into a Parquet file for efficient downstream analysis.

### **Supplementary Note 2: Feature Catalog**

DeepLogBot computes behavioral features per location, organized into several categories. The **36 features used by the fusion meta-learner** are marked in **Table** **1**; these are the features passed as input to the *GradientBoosting* classifier. Additional features are computed during extraction but serve primarily as intermediate signals for seed selection or diagnostics, *not* as inputs to the meta-learner. **Table** **1** provides the complete catalog.

**Table 1**: Complete feature catalog organized by category.

| **Feature** | **Description** |
| --- | --- |
| **Feature** | **Description** |
| ***Basic Activity (25 features)*** | |
| unique_users | Number of distinct user identifiers |
| downloads_per_user | Total downloads / unique users |
| total_downloads | Aggregate download count |
| projects_per_user | Unique datasets / unique users |
| avg_users_per_hour | Average unique users across active hours |
| max_users_per_hour | Peak unique users in any single hour |
| user_cv | Coefficient of variation of users per hour |
| users_per_active_hour | Users per hour (only counting active hours) |
| hourly_download_std | Standard deviation of hourly download counts |
| peak_hour_concentration | Fraction of downloads in the busiest hour |
| working_hours_ratio | Fraction during 9AM–6PM local time |
| hourly_entropy | Shannon entropy of hourly distribution |
| night_activity_ratio | Fraction during 10PM–6AM |
| yearly_entropy | Distribution uniformity across years |
| peak_year_concentration | Max year’s fraction of all downloads |
| years_span | Number of years with activity |
| downloads_per_year | Average annual download count |
| year_over_year_cv | CV of yearly download counts |
| fraction_latest_year | Latest year’s share of total |
| is_new_location | Binary: only active in latest year |
| spike_ratio | Latest year / historical average |
| unique_projects | Number of distinct datasets accessed |
| top_project_concentration | Fraction of downloads from the single most-downloaded project |
| top3_project_concentration | Fraction of downloads from the three most-downloaded projects |
| project_hhi | Herfindahl-Hirschman Index of per-project download shares |
| ***Advanced Behavioral (15 features)*** | |
| burst_pattern_score | Concentration of downloads in short time windows |
| circadian_rhythm_deviation | Deviation from typical human circadian patterns |
| user_coordination_score | Synchronized activity across user IDs |
| hourly_cv_burst | CV of hourly counts (burst detection) |
| spike_intensity | Magnitude of download spikes |
| user_peak_ratio | Peak-hour users / average users |
| night_ratio_advanced | Refined night activity metric |
| work_ratio_advanced | Refined working hours metric |
| evening_ratio | 6PM–10PM activity fraction |
| morning_ratio | 6AM–9AM activity fraction |
| user_coordination_std | Variability in coordination patterns |
| avg_concurrent_users | Average concurrent active users |
| max_concurrent_users | Peak concurrent users |
| is_bursty_advanced | Binary: exhibits burst patterns |
| is_nocturnal | Binary: predominantly night activity |
| ***Bot Interaction (6 features)*** | |
| dl_user_per_log_users | Downloads per user normalized by log(users) |
| user_scarcity_score | Inverse user density measure |
| download_concentration | Gini of download distribution across users |
| temporal_irregularity | Non-uniformity of temporal access patterns |
| bot_composite_score | Weighted combination of bot indicators |
| anomaly_dl_interaction | Anomaly score $\times$ download concentration |
| ***Bot Signature (8 features)*** | |
| request_velocity | Downloads per active day |
| access_regularity | Regularity of access intervals |
| ua_per_user | User agent diversity per user |
| ip_concentration | IP address concentration |
| session_anomaly | Deviation from normal session patterns |
| request_pattern_anomaly | Unusual request sequences |
| weekend_weekday_imbalance | Weekend vs weekday activity ratio |
| is_high_velocity | Binary: extremely high request rate |
| ***Discriminative (17 features)*** | |
| file_exploration_score | Breadth of file access patterns |
| file_mirroring_score | Consistency with mirroring behavior |
| file_entropy | Shannon entropy of file access distribution |
| bot_farm_score | User homogeneity (coordinated fakes) |
| user_authenticity_score | Diversity of user behavior patterns |
| user_homogeneity_score | Similarity across users at location |
| geographic_stability | IP geographic consistency |
| version_concentration | Concentration in file versions |
| targets_latest_only | Binary: only accesses latest versions |
| unique_versions | Number of distinct file versions |
| lifespan_days | Duration of activity |
| activity_density | Active days / total lifespan |
| persistence_score | Long-term consistent access pattern |
| malicious_bot_score | Composite malicious indicator |
| legitimate_auto­mation_score | Composite legitimate indicator |
| bot_vs_legit­imate_score | Differential (malicious $-$ legitimate) |
| is_likely_malicious | Binary: likely malicious automation |
| ***Time Series (23 features)*** | |
| *Outburst Detection (6 features)* | |
| outburst_count | Number of spikes ($>2$ standard deviations) |
| outburst_intensity | Average spike magnitude |
| max_outburst_zscore | Highest Z-score across time windows |
| outburst_ratio | Fraction of activity in outbursts |
| time_since_last_outburst | Recency of latest spike |
| longest_outburst_streak | Max consecutive high-activity periods |
| ***Periodicity Detection (4 features)*** | |
| weekly_autocorr | Autocorrelation at 7-day lag |
| dominant_period_days | Most significant period (FFT) |
| periodicity_strength | Strength of dominant period |
| period_regularity | Consistency of the period |
| ***Trend Analysis (5 features)*** | |
| trend_slope | Linear trend direction (normalized) |
| trend_strength | $R^{2}$ of linear fit |
| trend_acceleration | Second derivative |
| detrended_volatility | Volatility after removing trend |
| trend_direction | Categorical ($-1$, $0$, $+1$) |
| ***Recency-Weighted (4 features)*** | |
| recent_activity_ratio | Recent 30 days vs historical average |
| recent_volatility_ratio | Recent CV vs historical CV |
| recency_concentration | Fraction in last 30 days |
| momentum_score | Exponentially-weighted trend |
| ***Bot Signature Temporal (3 features)*** | |
| autocorrelation_lag1 | Day-to-day correlation |
| circadian_deviation | Distance from human circadian pattern |
| request_timing_entropy | Entropy of request timing |

### **Supplementary Note 3: Algorithm Details**

**Rule-Based Method:** The rule-based method uses threshold patterns defined in a YAML configuration file. Classification proceeds in two stages to assign each location to one of three categories: **bot**, **hub** (legitimate automation), or **user**:

1. **Stage 1 (User vs. Automated):** User patterns match on working_hours_ratio $\geq0.4$ and regularity_score $\leq0.6$, or interval_cv $\geq0.7$, or unique_users $<50$ with moderate activity. Automated patterns match on regularity_score $\geq0.7$, or night_activity_ratio $\geq0.35$ with low working hours, or user_coordination_score $\geq0.6$ with many users.
2. **Stage 2 (Bot vs. Hub):** Among automated locations, bot patterns include many-users-low-downloads (unique_users $\geq1000$, downloads_per_user $\leq50$), coordinated activity (coordination_score $\geq0.7$, authenticity_score $\leq0.4$), and suspicious timing (night_activity_ratio $\geq0.5$, working_hours_ratio $\leq0.2$). Hub patterns match on mirror-like behavior (downloads_per_user $\geq500$, unique_users $\leq100$) or CI/CD patterns (users $\leq10$, regularity $\geq0.7$).
3. **Stage 3 (Hub Protection):** The same structural hub protection rules used by the deep pipeline are applied as a post-classification safety net, overriding bot labels for locations with definitive institutional patterns (e.g., high downloads per user over multiple years, protocol-based detection via Aspera/Globus).

**Deep Architecture Method:** The deep method implements a four-phase semi-supervised pipeline (Seed Selection, LLM Seed Refinement, Deep learning algorithm, Hub Protection & Finalization). This section provides a detailed description of each phase, including the specific thresholds, design rationale, known edge cases, and failure modes.

1. **Phase 1** (**Seed Selection**): Seed selection identifies high-confidence training examples for each category using structural and behavioral heuristics. The seeds are *not* the final classification; they serve only as labeled training data for the meta-learner. Each seed receives a confidence weight (0–1) that influences its importance during gradient-boosted training.
   a. **Minimum volume filter**: All seeds require a minimum of 20 total downloads (MIN_SEED_DOWNLOADS $=20$). Below this threshold, behavioral features (e.g., hourly entropy, working hours ratio) become unreliable because they are computed from too few events. This filter is critical: without it, the training set would include noisy locations where a handful of downloads happened to fall at night or during working hours purely by chance, degrading meta-learner performance.
   b. **User seeds** (3-tier system).
      i. **Tier A - Individual researchers** (confidence 1.0): $\leq$10 users, $\leq$5 downloads/user, 20–200 total downloads, working hours ratio $>$0.3, night activity $<$0.5, $\geq$2 years span.
      ii. **Tier B - Active researchers** (confidence 0.7): $\leq$50 users, $\leq$10 downloads/user, 20–1000 total downloads, working hours ratio $>$0.25, hourly entropy $>$1.5, burst pattern score $<$0.5.
      iii. **Tier C - Research groups** (confidence 0.4): $\leq$200 users, $\leq$20 downloads/user, $\geq$20 total downloads, user coordination score $<$0.3, protocol legitimacy $>$0.3. Additional safeguards exclude locations active only in the latest year with $>$50 users (likely distributed bot farms appearing as “new” research groups) and single-year locations with $>$30 users (insufficient history to confirm user behavior).
   c. **Bot seeds** (6 complementary signals):
      i. **Bot farm**: $>$5,000 users, $<$50 downloads/user. The classic distributed bot-farm signature: many distinct user identifiers each performing minimal activity, consistent with rotating IP addresses or user agents.
      ii. **Distributed bot network**: 500–5,000 users, 2–15 downloads/user, working hours ratio $<$0.38, fraction of downloads in the latest year $>$0.8. Captures the “long tail” of bot locations with moderate user counts but strong temporal signals (no circadian rhythm, sudden recent appearance).
      iii. **Nocturnal**: Night activity ratio $>$0.8, working hours ratio $<$0.1. Extreme nocturnal activity with essentially no daytime presence, inconsistent with any plausible human usage pattern regardless of time zone.
      iv. **Coordinated**: $>$10,000 users, $<$20 downloads/user. Massive-scale coordinated access characteristic of large botnets or web crawlers.
      v. **Scraper**: $>$15,000 unique projects accessed. Locations that systematically crawl the entire PRIDE catalog, accessing far more datasets than any researcher or institution would.
      vi. **Year-over-year explosion**: Spike ratio $>$50$\times$ compared to previous years, $>$200 users, $>$95% of activity in the latest year. Captures locations that experience sudden, extreme surges in activity, typically indicating the emergence of a new automated process rather than gradual growth.
      vii. **Hub-like exclusion from bot seeds**: Locations with downloads/user $>$200 sustained over $\geq$3 years are excluded from bot seeds regardless of other signals. This prevents institutional mirrors from contaminating the bot training set; a mirror may have thousands of users and uniform temporal patterns, but the key distinguishing feature is high downloads per user (institutional users download hundreds of files each, while bot “users” download 3–15). This exclusion is critical for preventing the meta-learner from learning to associate high download volume with bot behavior.
   d. **Hub seeds** (2 structural patterns).
      i. **Small mirrors**: Downloads/user $>$500, $<$100 users. Classic institutional mirrors or automated reanalysis pipelines operated by a small team.
      ii. **Institutional hubs**: Downloads/user $>$200, $\leq$1,000 users, $\geq$3 years span. Larger institutions with sustained high-volume access over multiple years, consistent with established bioinformatics centers.

Hub seeds exclude nocturnal-dominant locations (working hours ratio $<$0.1 and night activity $>$0.7) to avoid capturing bot locations that happen to have high downloads/user. Confidence is set to 0.8 (base), boosted to 0.95 for protocol-verified hubs (Aspera ratio $>$0.3 or Globus ratio $>$0.1) and 0.85 for long-running hubs ($\geq$4 years).

**Seed overlap resolution:** Seed sets are resolved with strict priority ordering: **hub** $>$ **bot** $>$ **user**. If a location qualifies as both a hub seed and a bot seed, it is retained only as a hub seed and removed from the bot set. Similarly, bot–user overlaps are resolved in favor of the bot.

2. **Phase 2 (LLM Seed Refinement):** The blind multi-LLM consensus annotation and seed injection procedure is described in detail in **Supplementary Note 4**. This phase produces 934 validated consensus labels from two independent LLM annotators, of which 625 are injected as high-confidence training seeds.
3. **Phase 3 (Fusion Meta-Learner)**: A GradientBoostingClassifier (200 estimators, max depth 5, learning rate 0.1, subsample 0.8, min samples per leaf 10) is trained on the seed sets with confidence-weighted samples. Features are standardized using StandardScaler before training to ensure features on different scales contribute equally.
4. **Phase 4 (Hub Protection & Finalization):** Structural hub protection rules act as a post-classification safety net, overriding the meta-learner’s output for locations with definitive institutional patterns. This phase addresses a key architectural asymmetry: **the cost of misclassifying a hub as a bot is much higher than the reverse.** Labeling an institutional hub as a bot permanently excludes legitimate scientific infrastructure from download statistics, whereas a bot misclassified as a hub inflates hub counts but does not distort user-level analyses. The protection rules operate on two structural signals from the config file:
   a. **Protocol-based hub detection**: Aspera ratio $>$0.3 or Globus ratio $>$0.1. These high-performance bulk-transfer protocols are almost exclusively used by institutional infrastructure.
   b. **Extreme mirrors**: Downloads/user $>$500, $\leq$200 users. Classic mirror pattern with very high per-user download intensity and few distinct users.

### **Supplementary Note 4: Ground Truth Construction**

Initial ground truth labels were assigned using high-confidence heuristic criteria applied to a 1-million record sample (**Table 2**):

**Table 2**: Ground truth label criteria and counts. Subtypes are applied hierarchically (each excludes locations matched by prior subtypes within the same category); per-subtype counts are not listed as they depend on evaluation order; totals per category are the relevant quantities.

| Label | Subtype | Count | Criteria |
| --- | --- | --- | --- |
| Bot | Ground truth bot | – | $\geq$10K users, $\leq$10 DL/user |
|  | Large-scale bot | – | $\geq$5K users, $\leq$25 DL/user |
|  | Bot farm | – | $\geq$1K users, $\leq$50 DL/user, work ratio $\leq$0.3 |
| 2-4 | **Total** | **88** |  |
| Hub | Mirror | – | $\leq$5 users, $\geq$1K DL/user |
|  | Institutional hub | – | $\leq$20 users, $\geq$500 DL/user |
|  | Research hub | – | 10–200 users, $\geq$200 DL/user, $\geq$100K total, work ratio $\geq$0.2 |
| 2-4 | **Total** | **44** |  |
| User | Individual user | – | $\leq$3 users, $\leq$20 DL/user, work ratio $\geq$0.4 |
|  | Research group | – | 3–30 users, 5–100 DL/user, work ratio $\geq$0.35 |
|  | Casual user | – | $\leq$5 users, $\leq$50 DL/user, night ratio $\leq$0.3 |
| 2-4 | **Total** | **1,279** |  |
| Uncertain | – | 18,634 | Excluded from benchmark |

**Blind Multi-LLM Independent Validation:** To address the circularity inherent in evaluating a semi-supervised classifier against its own labels, we constructed an independent validation set using blind multi-LLM consensus annotation. Critically, neither LLM was shown DeepLogBot’s classification labels or confidence scores, eliminating anchoring bias. We sampled 1,153 locations from the full 71,133-location dataset using 20 stratified zones designed to cover the full feature space, including both easy and difficult classification regions.

Two LLMs annotated all 1,153 locations independently and without access to any classifier outputs. Each LLM received only: (i) the city, country, and geographic coordinates; (ii) 14 behavioral features (user count, downloads per user, working hours ratio, night activity, protocol usage, temporal span, etc.); and (iii) geographic research context enriched via EuropePMC publication counts. The classifier’s labels, confidence scores, and all derived classification columns were explicitly excluded to prevent anchoring bias.

The two annotators were:

1. **Claude Opus 4.6** (Anthropic): Run via Claude Code subagents in 24 parallel batches of $\sim$50 locations each, with no shared context between batches.
2. **Qwen3-30B-A3B** (Alibaba): Run locally via Ollama with temperature 0.1 for consistency, processing each location sequentially with no access to other annotations.

Both LLMs used the same structured prompt defining three categories (bot, hub, user) with explicit feature interpretation guidelines. The key discriminator was downloads per user: bots typically show 3–15 DL/user, hubs $>$200 DL/user, and users fall in between with research-consistent patterns.

### **Supplementary Note 5: Rule-based algorithm vs deep-learning**

We ran the rule-based classifier on the complete 159.3M-record dataset and evaluated both methods against the 934 blind multi-LLM consensus labels (**Table 3**).

**Table 3**: Full-dataset classification comparison: Rules vs Deep. Location counts reflect full-dataset classification; “User locs” includes both classified user locations and locations with insufficient evidence ($<$3 downloads). Accuracy and Macro F1 are measured against all 934 blind multi-LLM consensus labels; of these, 625 were used as training labels for the Deep method.

| **Method** | **Bot locs** | **Hub locs** | **User locs** | **Accuracy** | **Macro F1** |
| --- | --- | --- | --- | --- | --- |
| Rules | 22,981 | 714 | 47,438 | 59.7% | 0.592 |
| Deep | 27,063 | 249 | 43,821 | 92.2% | 0.909 |

The rule-based method identifies fewer bot locations than the deep pipeline (22,981 vs. 27,063). This underdetection arises from two main issues: (i) fixed thresholds only flag locations that clearly exceed preset criteria (e.g., >1,000 users, ≤50 DL/user), missing distributed bots with moderate activity; and (ii) any location not matching a predefined automated pattern is labeled as a user, even when its behavior suggests otherwise. In contrast, the deep pipeline jointly evaluates all 36 behavioral features and leverages gold-standard training labels, allowing it to detect bot patterns missed by single-threshold rules.

**Table 4**: Per-class precision, recall, and F1 for each method against 934 blind multi-LLM consensus labels.

| Method | Class | Precision | Recall | F1 |
| --- | --- | --- | --- | --- |
| Rules | Bot | 0.810 | 0.589 | 0.682 |
|  | Hub | 0.653 | 0.858 | 0.742 |
|  | User | 0.287 | 0.454 | 0.352 |
| Deep | Bot | 1.000 | 0.975 | 0.987 |
|  | Hub | 0.687 | 1.000 | 0.815 |
|  | User | 1.000 | 0.778 | 0.875 |

The rule-based method achieves reasonable bot precision (0.810) but the lowest user F1 (0.352), confirming that static thresholds cannot separate research-city user traffic from automated patterns.

### **Supplementary Note 6: Bot Removal Analysis**

**Full-Dataset Classification:** The semi-supervised classification pipeline was applied to the complete dataset of locations aggregated from 159.3 million download records. The classification results are summarized in Figure 3. The asymmetry between location counts and download volumes is striking bots generate far more traffic per location on average, while institutional hubs, though comprising a small fraction of locations, account for a substantial share of download volume.

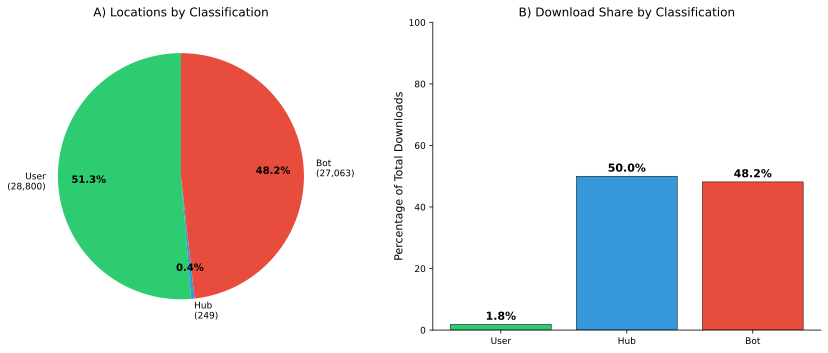

**Figure 3**: Full-dataset classification results. (A) Location distribution by classification category. (B) Download distribution: bots generate the majority of all traffic.

### **Supplementary Note 7: Extended Usage Analysis**

**Monthly Download Trends**: Monthly download patterns (after bot removal) reveal temporal structure within the yearly trends (**Figure 4**). Activity shows seasonal variation with increased downloads during the academic year and a notable surge in early 2025, likely reflecting both genuine growth and the inclusion of January 2025 data.

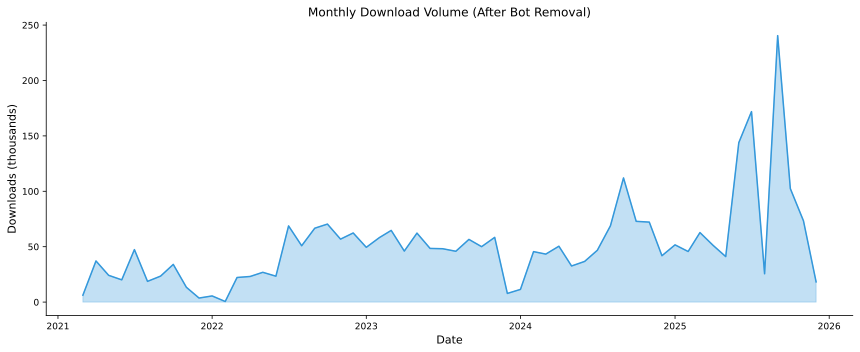

**Figure 4**: Monthly download volume (after bot removal), showing seasonal patterns and overall growth.

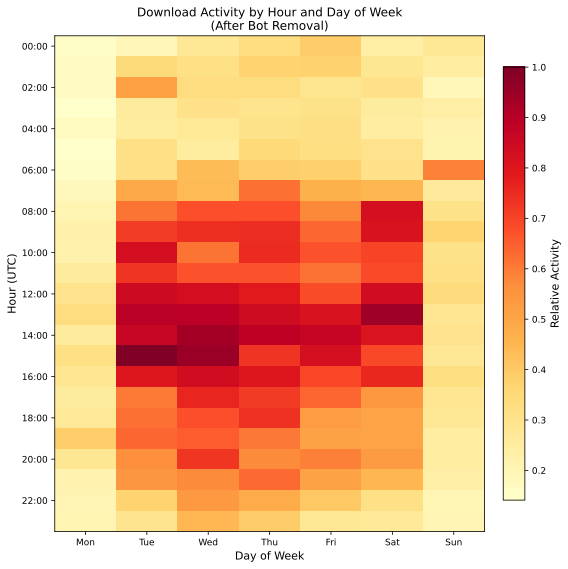

**Figure 5**: Download activity heatmap by hour (UTC) and day of week, after bot removal. The circadian pattern confirms genuine human usage.

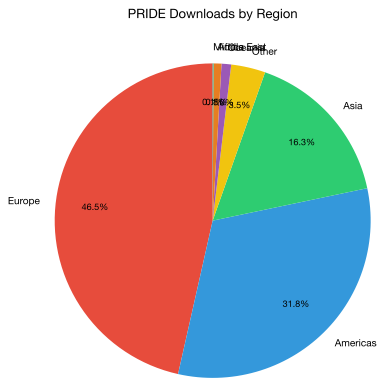

**Figure 6**: PRIDE downloads by world region (after bot removal, 2021–2025).

**pridepy adoption:** To facilitate the adoption of high-performance download protocols, the PRIDE team released pridepy (<https://github.com/PRIDE-Archive/pridepy>). The package was published on PyPI in March 2025. **Figure 7** shows the monthly download trend throughout 2025, with a total of 6,504 installations. Notably, downloads surged in October 2025 (1,111 installations), coinciding with the rapid growth of Aspera-based transfers observed in the PRIDE download logs.

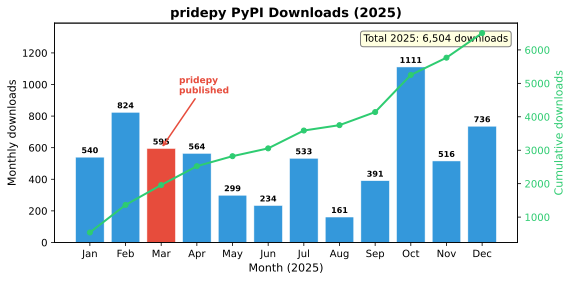

**Figure 7:** Monthly PyPI download statistics for pridepy throughout 2025. The red bar marks the publication month (March 2025). The green line shows cumulative downloads. Data source: pypistats.org (August–December 2025).

**Top Downloaded Datasets: Figure 8** shows the top 20 PRIDE datasets ranked by hub download volume, illustrating which datasets are most targeted by institutional mirrors and reanalysis infrastructure. The most downloaded dataset by hubs is **PXD012988** (1.77M hub downloads), followed by **PXD012987** (1.52M) and **PXD012162** (1.09M). These hub-dominated datasets differ substantially from the user-download rankings (**Figure 4B** in the main text), where PXD010154 (31.4K user downloads from 43 countries) leads, followed by PXD001819 (23.3K) and PXD000561 (20.5K from 60 countries). This divergence confirms that hub traffic reflects institutional mirroring priorities rather than individual researcher interest and underscores the importance of separating traffic categories when assessing dataset impact.

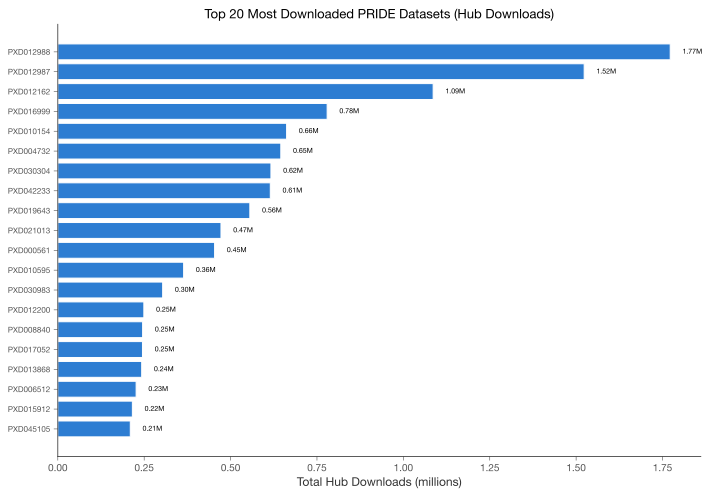

**Figure 8**: Top 20 most downloaded PRIDE datasets by hub download volume, showing the datasets most targeted by institutional mirrors and reanalysis infrastructure.

**File-Level Download Distribution**: At the individual file level, downloads follow a log-normal distribution (**Figure 9**), with most files receiving between 3 and 30 downloads. A small number of files exceed 1,000 downloads, representing benchmark datasets and popular reference proteomes.

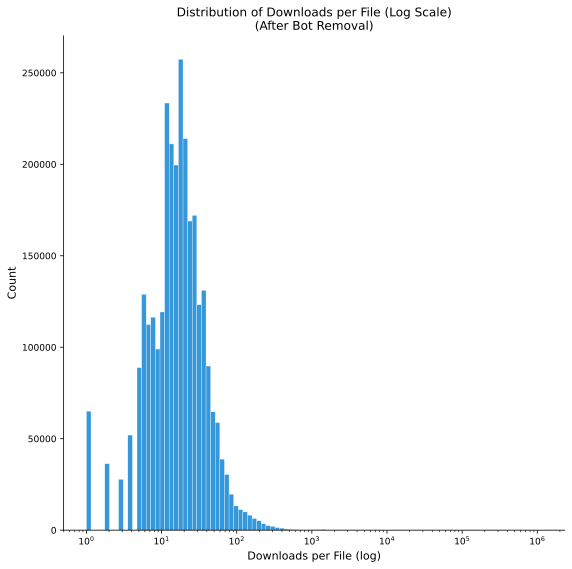

**Figure 9**: Distribution of downloads per file (log scale). The majority of files receive between 3-30 downloads.

### **Supplementary Note 8: Limitations**

**Semi-supervised label circularity:** The seed labels used to train the fusion meta-learner are derived from heuristic rules rather than independently verified ground truth. Although the six bot signals and structural hub criteria capture well-understood traffic patterns, any systematic bias in the seeds propagates through the learned model. To mitigate this, we construct a stratified validation set of 1,153 locations spanning 20 feature-space zones, annotated blindly by two independent LLMs without access to classifier outputs, yielding 934 consensus labels reviewed by a domain expert.

***User identity resolution:*** A “unique user” is defined as a distinct anonymized IP hash. This proxy is imperfect in two directions: (i) multiple individuals behind a single NAT gateway or institutional proxy appear as one user, deflating user counts; and (ii) a single individual using multiple networks (e.g., VPN, home vs. office) appears as multiple users, inflating counts. Consequently, the “downloads per user” metric, central to both hub protection and bot-farm detection, may be distorted for locations with heavy proxy usage.

**Geographic location granularity:** All users from the same geolocated coordinate are grouped into a single location profile. In large metropolitan areas, this conflates distinct institutions and user populations (e.g., Beijing), potentially masking mixed user/bot behavior. Conversely, users at the same institution on different subnets may map to different coordinates, splitting what is logically one location.

**IP geolocation accuracy**: Geographic attribution relies on MaxMind GeoIP databases, whose accuracy varies by region. Locations in sub-Saharan Africa, Central Asia, and small island states may be misattributed, and cloud-hosted downloads (AWS, Google Cloud) are mapped to data-center locations rather than the researcher’s actual location.

**Hub protection threshold sensitivity**: The hub protection rules use fixed thresholds (e.g., downloads per user $>200$, years span $\geq3$) chosen based on empirical inspection of known research institutions. These thresholds may be too permissive for domains with different download patterns (e.g., a repository where legitimate per-user download volumes are lower) or too restrictive if large-scale automation becomes more prevalent.
